# Targeting Astrocytic Stat3 Reveals Context-Dependent Modulation of Prion Disease

**DOI:** 10.64898/2026.08.15.745015

**Authors:** Natallia Makarava, Narayan P. Pandit, Olga Mychko, Kara Molesworth, Tarek Safadi, Olga Bocharova, Ilia V. Baskakov

**Affiliations:** Department of Neurobiology, University of Maryland School of Medicine, Baltimore, Maryland, USA; Department of Anesthesiology and Center for Shock, Trauma and Anesthesiology Research (STAR), University of Maryland School of Medicine, Baltimore, Maryland, USA

**Keywords:** prion diseases, prions, Stat3, reactive astrocytes, neuroinflammation, prion strains, sexual dimorphism, GFAP, Vimentin

## Abstract

Reactive astrogliosis is a prominent feature of prion diseases, yet the molecular mechanisms regulating astrocyte activation and their contribution to disease progression remain poorly understood. Signal transducer and activator of transcription 3 (Stat3) is a master regulator of reactive astrocytes in numerous neurological disorders, but its role in prion disease has not been established. Here, we investigated the contribution of astrocytic Stat3 signaling to prion pathogenesis using an inducible astrocyte-specific Stat3 knockout mouse model. Stat3 expression was elevated across multiple neuroinflammatory conditions but was most strongly induced during prion disease. Among four mouse-adapted prion strains (ME7, RML, 22L, and SSLOW), the magnitude of Stat3 activation closely paralleled the severity of neuroinflammation. Astrocyte-specific Stat3 deletion was evaluated in mice infected with either the highly inflammatory SSLOW strain or the less inflammatory 22L strain. Stat3 deletion had no detectable effect on disease progression in SSLOW-infected mice but modestly delayed disease onset and behavioral decline in male mice infected with the 22L strain, particularly when knockout was induced before prion inoculation. Despite its limited effect on survival, astrocyte-specific Stat3 deletion consistently attenuated astrocyte reactivity, as evidenced by reduced vimentin expression, delayed cortical GFAP induction, and lower GFAP expression in recombined astrocytes at the single-cell level, demonstrating a cell-autonomous role for Stat3 in promoting reactive astrogliosis. In contrast, PrP^Sc^ accumulation and overall microglial activation remained unchanged, indicating that astrocytic Stat3 signaling is dispensable for prion replication and does not substantially influence the global microglial response. Tamoxifen-induced recombination occurred in only 40-70% of astrocytes, resulting in partial and region-dependent Stat3 deletion that likely underestimated the impact of astrocytic Stat3 loss. Together, these findings identify Stat3 as an important regulator of astrocyte reactivity during prion disease but demonstrate that its contribution to disease progression is limited and highly context-dependent, varying with the inflammatory milieu, timing of pathway inhibition, and biological sex. Our results highlight the redundancy of inflammatory signaling networks driving chronic prion neurodegeneration and suggest that targeting astrocytic Stat3 alone is unlikely to substantially alter disease progression.

## Introduction

Prion diseases are fatal neurodegenerative disorders characterized by the conformational conversion of the cellular prion protein (PrP^C^) into a misfolded, self-propagating isoform (PrP^Sc^) [1–3]. Progressive accumulation of PrP^Sc^ triggers synaptic dysfunction, neuronal loss, spongiform degeneration, and robust activation of glial cells throughout the central nervous system [4–11]. While the mechanisms governing prion replication have been extensively investigated, increasing evidence indicates that disease progression is profoundly influenced by the host glial response [12–22]. Astrocytes and microglia undergo extensive phenotypic remodeling during prion disease [6, 7, 13, 16, 23–26], yet the molecular pathways regulating these responses and their contributions to neurodegeneration remain poorly understood.

Astrocytes are highly heterogeneous cells that perform numerous homeostatic functions, including regulation of neurotransmitter metabolism, maintenance of extracellular ion balance, metabolic support of neurons, preservation of blood-brain barrier integrity, and modulation of synaptic activity [27–29]. In response to central nervous system injury or disease, astrocytes undergo a process broadly referred to as reactive astrogliosis, characterized by cellular hypertrophy, increased expression of intermediate filament proteins such as glial fibrillary acidic protein (GFAP) and vimentin, transcriptional reprogramming, and functional changes that vary according to disease context [23, 30–34]. Rather than representing a single activation state, reactive astrocytes encompass a spectrum of phenotypes ranging from neuroprotective responses that promote tissue repair to maladaptive states that exacerbate neuroinflammation and neuronal dysfunction [35–39]. Defining the molecular mechanisms that govern these diverse astrocytic states has become a major objective in neurodegeneration research.

Among the signaling pathways implicated in reactive astrogliosis, the Janus kinase (Jak2)/signal transducer and activator of transcription 3 (Stat3) pathway has emerged as a central regulator [40–42]. Stat3 is activated downstream of multiple cytokines and growth factors, including members of the IL-6 cytokine family, leukemia inhibitory factor (LIF), ciliary neurotrophic factor (CNTF), and other inflammatory mediators [43–45]. Following phosphorylation, Stat3 dimerizes and translocates to the nucleus, where it regulates transcription of genes involved in astrocyte hypertrophy, intermediate filament expression, cytokine production, extracellular matrix remodeling, and tissue repair. Genetic studies have established Stat3 as a master regulator of astrocyte reactivity following acute and chronic CNS conditions [41, 46–52]. Astrocyte-specific deletion of Stat3 markedly attenuated reactive astrogliosis after spinal cord injury, and altered inflammatory responses [45, 50–52]. These studies demonstrate that Stat3-dependent transcription is essential for the structural and functional remodeling of reactive astrocytes. In chronic neurodegenerative disorders, the role of astrocytic Stat3 functions appear to be highly context dependent. Depletion of Stat3 has been reported to reduce astrocyte reactivity, suppressed microglia activation, reduced amyloid deposition, restored synaptic deficits and improve cognitive performance in mouse models of Alzhemier’s disease [46–48]. Contrary, in Huntington’s disease, activation of astrocytic Jak2-Stat3 pathway promotes proteostasis and enhance clearance of mutant Huntingtin in neurons. These findings underscore the complexity of Stat3-mediated astrocyte responses and suggest that reactive astrogliosis cannot be universally classified as either protective or harmful (reviewed in [38–40, 53]).

Prion diseases exhibit one of the most robust astrocytic responses among neurodegenerative disorders [37]. Astrocyte activation occurs early during disease progression, preceding overt neuronal loss, and becomes increasingly prominent as PrP^Sc^ accumulates throughout the brain [16, 24, 32–34]. Transcriptomic analyses have demonstrated substantial induction of inflammatory and stress-response pathways in astrocytes during prion infection, including activation of genes associated with cytokine signaling, complement cascade components, lysosomal pathways, and extracellular matrix remodeling [16, 33, 34]. Inverse correlation between the extent of astrocyte reactivity and the incubation period for prion disease in mice infected with a range of prion strain suggests that astrocyte reactivity contribute to disease progression [16]. Supporting this hypothesis, reactive astrocytes isolated from prion-infected animals were shown to exert harmful effects on primary neuronal cultures, reducing spine size and density and impairing neuronal growth and synaptic integrity [54]. Likewise, reactive astrocytes derived from prion-infected mice induced a disease-associated phenotype in endothelial cells from non-infected adult mice [55]. Retraction of astrocytic endfeet from blood vessels and loss of BBB integrity observed at late preclinical stage of the disease also point at involvement of reactive astrocytes in disease pathogenesis [16, 55].

Previous studies have identified Stat3 signaling as a prominent feature of reactive astrocytes during prion disease [56–58]. However, the functional significance of astrocytic Stat3 signaling in prion disease has remained unclear. Because IL-6 family cytokines and oncostatin M, potent activators of Stat3, are elevated during prion disease, we proposed that sustained activation of this pathway is thought to represent a major driver of astrocyte reactivity [57].

In the present study, we investigated the functional role of astrocytic Stat3 signaling in prion disease using an inducible astrocyte-specific Stat3 knockout mouse model. We demonstrate that Stat3 is robustly activated in reactive astrocytes across multiple prion strains, with the magnitude of activation correlating with the severity of neuroinflammation. Unexpectedly, astrocyte-specific Stat3 deletion exerted only a modest effect on disease progression, delaying clinical onset primarily in male mice infected with the less inflammatory 22L strain, while having no detectable effect in the highly inflammatory SSLOW strain. Despite its limited impact on disease progression, Stat3 deletion attenuated astrocyte reactivity by delaying the development of the reactive phenotype without affecting PrP^Sc^ accumulation or microglial activation. Finally, we show that the incomplete and region-dependent efficiency of tamoxifen-induced recombination resulted in only partial astrocyte-specific Stat3 deletion, a factor that likely contributed to the modest phenotype observed. Together, these findings establish Stat3 as an important regulator of astrocyte reactivity during prion disease while suggesting that astrocytic Stat3 signaling plays a relatively limited role in determining overall disease progression.

## Methods

### Animals

Aldh1l1-CreERT-Stat3-floxP mouse model was provided by Dr. Sofroniew [59]. For generating mice for astrocyte-specific Stat3 knock-out (Aldh1l1-CreERT)^+/-^(Stat3-floxP)^+/+^ and littermate controls (Stat3-floxP)^+/+^ denoted as Cre^+/−^ and Cre^−/−^, respectively, mice were bred as previously described [57]. C57BL/6J wild type mice (Veterinary Resources, University of Maryland, Baltimore, Maryland, USA) were used as an additional control (WT). Mouse models of neurological conditions used for NanoString analysis of Stat3 expression were described previously [37].

A 20 mg/mL stock solution of tamoxifen (#T5648, Sigma) was prepared by dissolving tamoxifen in one part of 100% ethanol, followed by the addition of nine parts of corn oil prewarmed to 37°C. The mixture was continuously shaken at 37°C until fully dissolved. To induce conditional Stat3 knockout, male and female (Aldh1l1-CreERT)^+/−^(Stat3-floxP)^+/+^ and their (Stat3-floxP)^+/+^ littermates were subjected to subcutaneous (SQ) tamoxifen injections, 100 mg/kg body weight, once every 24 hours for a total of five consecutive days. Depending on experimental setup, mice received tamoxifen injections at 4 – 9 weeks of age (before infection with prions), or 10 – 14 weeks after prion infection, as described.

### Prion infection and disease monitoring

10% (w/v) brain homogenates from mice terminally ill with prion disease were prepared in PBS, pH 7.4, using glass/Teflon homogenizers attached to a cordless 12 V compact drill, and stored at -80°C [60]. Immediately before inoculation, each homogenate was further dispersed by 30 seconds of indirect sonication at approximately 200 watts in a microplate horn of a sonicator (Qsonica, Newtown, CT) and diluted in PBS, as needed. Except where mentioned otherwise, male and female mice were intracerebrally (IC) inoculated with 20 µl volume of 10% SSLOW or 1% 22L mouse-adapted prion strains in PBS under 3% isoflurane anesthesia. In one set of experiments, five- to eight-week-old Cre^+/−^, Cre^−/−^, and WT mice were first subjected to IC inoculation with prions, and then Cre^+/−^ and Cre^−/−^ mice were treated with tamoxifen 10 - 14 weeks later, as described above. In an alternative experimental design, four- to nine-week-old Cre^+/−^, Cre^−/−^ mice were treated with tamoxifen first, and then IC inoculated with prions 4 weeks later, together with their WT counterparts. Mice were regularly monitored for clinical signs, which included clasping hind legs, difficulty walking, abnormal gait, nesting problems, and weight loss.

Starting at the preclinical stage, mice were tested once per week in an elevated plus maze (EPM). In each session, a mouse was placed at the center of EPM and given 5 minutes to explore the maze. All movements were captured using video recording, and total timing spent in open arms, closed arms, or the center of the maze was quantified using ANY-maze software (version 6.33). After the first training sessions, mice naturally acquire a strong preference for the closed arms, i.e., avoiding open arms during the sessions that follow the training session until the clinical onset of the disease. The mice were considered terminal and euthanized when they were unable to rear and/or lost 20% of their weight.

### Antibodies

Primary antibodies used for immunofluorescence and immunoblotting were as follows: chicken polyclonal anti-GFAP (#AB5541, Millipore Sigma, Burlington, MA); rabbit monoclonal anti-GFAP, clone D1F4Q (#12389, Cell Signaling, Danvers, MA); rabbit monoclonal anti-Stat3, clone D1A5 (#8768, Cell Signaling); rabbit monoclonal anti-phospho-Stat3 (Tyr705), clone D3A7 (#9145, Cell Signaling); rabbit monoclonal anti-HA-tag, clone C29F4 (#3724, Cell Signaling); rabbit monoclonal anti-prion protein, clone 3D17 (#ZRB1268, Millipore Sigma); rabbit monoclonal anti-Vim, clone D21H3 (#5741, Cell Signaling); rabbit polyclonal anti-CD11b (ab128797, Abcam, Waltham, MA); rat monoclonal anti-Gal3, clone M3/38 (#sc-23938, Santa Cruz, Dallas, TX); mouse monoclonal anti-Tubb3, clone TUJ1 (#801201, BioLegend, San Diego, CA); rabbit polyclonal anti-IBA1 (#013-27691, FUJIFILM Wako Chemicals USA; Richmond, VA); goat polyclonal anti-IBA1 (#NB100-1028, Novus, Centennial, CO); mouse monoclonal anti-β-actin, clone AC-15 (#A5441, Sigma-Aldrich, Saint Luis, MO). The secondary antibodies for immunofluorescence were Alexa Fluor 488-, 546-, and 647-labeled (Thermo Fisher Scientific, Waltham, MA). The nuclei were stained with DAPI (#62248, Thermo Fisher Scientific).

### Immunofluorescence staining of mouse brains

Formalin-fixed brains (sagittal 3 mm slices) were treated for 1 hour in 96% formic acid before being embedded in paraffin using standard procedures; 4 μm sections produced with Leica RM2235 microtome (Leica Biosystems, Deer Park, IL) were mounted on Superfrost Plus Microscope slides (#22- 037-246, Thermo Fisher Scientific) and processed for immunohistochemistry according to standard protocols. To expose epitopes, slides were subjected to 20 minutes of hydrated autoclaving at 121°C in citrate buffer, pH 6.0, antigen retriever (#C9999, Sigma-Aldrich). For the detection of disease-associated PrP, an additional 3-minute treatment in concentrated formic acid was applied.

Autofluorescence Eliminator Reagent (#2160, Sigma-Aldrich) and Signal Enhancer (#11932, Cell Signaling Technology) were used on slides according to the original protocols to reduce background fluorescence. Images were collected using an inverted microscope Nikon Eclipse TE2000-U (Nikon Instruments Inc.) equipped with an illumination system X-cite 120 (EXFO Photonics Solutions Inc.) and a cooled 12-bit CoolSnap HQ CCD camera (Photometrics), or Leica Mica widefield microscope (Leica Microsystems, Boston, MA). Images were processed using ImageJ software (1.54p, NIH).

To measure GFAP intensity of HA^+^ and HA^−^ astrocytes, mouse brains collected at 91 dpi of 22L prion disease were co-immunostained for HA-tag and GFAP. The images acquired from cortex were subjected to automatic background subtraction and threshold to generate binary HA and GFAP images. Then, binary HA image was subtracted from binary GFAP image, and an area selection representing HA^−^ astrocytes was applied to the original GFAP channel to measure GFAP mean intensity of HA^−^ astrocytes. GFAP mean intensity of HA^+^ astrocytes was measured in an area selection created on the binary HA image.

To estimate the Cre recombinase efficiency, mouse brains collected at the terminal stage of the disease were co-immunostained for HA-tag and GFAP. The images acquired from cortex, hippocampus, thalamus and striatum were subjected to automatic background subtraction and threshold to generate binary HA and GFAP images. For each field of view, an ImageJ Image Calculator OR function was used to generate a combined binary image representing all astrocytes. Particle Analysis function of ImageJ was used for defining regions of interest (ROIs) representing individual astrocytes. Resulting ROIs were measured on binary HA images to estimate the number of HA^+^ and HA^−^ astrocytes.

### Western blot

For Western blots, 10% (w/v) brain homogenates (BH) were prepared using RIPA Lysis Buffer (#20-188, Millipore Sigma) supplemented with protease and phosphatase inhibitors (#5872S, Cell Signaling). To analyze brain-derived PrP^Sc^, BH aliquots were diluted with RIPA buffer to achieve 5% BH final concentration and treated with 20 µg/ml proteinase K (#P8107S, New England BioLabs, Ipswich, MA) in the presence of 50 mM Tris, pH 7.5, and 2% Sarcosyl, for 30 min at 37 °C. To analyze other proteins, BH was diluted with RIPA buffer to 1% and proteinase digestion was omitted. The resulting samples were supplemented with 4xSDS loading buffer and heated for 10 min in a boiling water bath before loading onto NuPAGE 12% Bis-Tris gels (#NP0341BOX, Thermo Fisher Scientific). Wet transfer onto PVDF membranes and probing of Western blots was done according to standard procedures. The signals were visualized by Immobilon Forte Western HRP Substrate (WBLUF0100, Millipore Sigma) or SuperSignal West pico PLUS Chemiluminescent Substrate (34577, Thermo Fisher Scientific) using Invitrogen iBright 1500 imager, and quantified with iBright Analysis software (Thermo Fisher Scientific). Intensity data were presented as normalized by actin, except for PrP^Sc^ Western blots treated with protease K.

### NanoString

NanoString experiments were previously described [16, 24]. Briefly, mice were euthanized, and their brains were immediately extracted and dissected to collect individual regions for RNA isolation using Trizol (Thermo Fisher Scientific, Waltham, MA, USA) and Aurum Total RNA Mini Kit (Bio-Rad, Hercules, CA, USA). An amount of 200 ng of total RNA was submitted to the Institute for Genome Sciences at the University of Maryland, School of Medicine, for RNA integrity check and subsequent analysis using an nCounter NanoString Neuroinflammation panel and custom nCounter Mouse Astrocyte Panel. Only samples with an RNA integrity number RIN > 7.2 were used for NanoString analysis. For each sample, the assessment of all target sequences was performed within a single tube, using uniquely coded hybridization probes designed by NanoString Thechnologies (Seattle, WA), enabling a reliable and reproducible assessment of the expression. All data passed quality control assessments for imaging, binding, positive control, or CodeSet content normalization. The analysis of data was performed using nSolver Analysis Software 4.0, including nCounter Advanced Analysis (version 2.0.115). Z-score transformation for heat maps was performed for genes.

### Statistics

Statistical analyses and plotting of the data were performed using GraphPad Prism software, versions 8.4.2–10.4.1 for Windows (GraphPad Software, San Diego, CA). Statistical comparison of two groups with normal distribution of data points and equal variances was made with an unpaired two-tailed t-test, except for a paired t-test comparison in Fig. 6D. Alternatively, non-parametric two-tailed Mann-Whitney test was used. In case of unequal variances, the comparison of two groups with normal distribution of data points was performed with an unpaired two-tailed t-test with Welch’s correction. For statistical comparison of more than two groups, one-way ANOVAs were followed by multiple comparison tests as mentioned in Figure Legends, and the resulting p-values were reported. For statistical analysis of data presented in Superplots, means ± SD were calculated using biological replicas, where n = the number of brains or animals analyzed.

## Results

### Stat3 is upregulated across multiple neurological conditions and is highly induced during prion disease

Bulk tissue mRNA analysis revealed increased Stat3 expression in the brains of mice across multiple neurological conditions associated with neuroinflammation, including aging, Alzheimer’s disease, traumatic brain injury, ischemia, and prion disease (Fig. 1A). Among these conditions, prion-infected mice exhibited the highest level of Stat3 upregulation (Fig. 1A).

**Figure 1.**
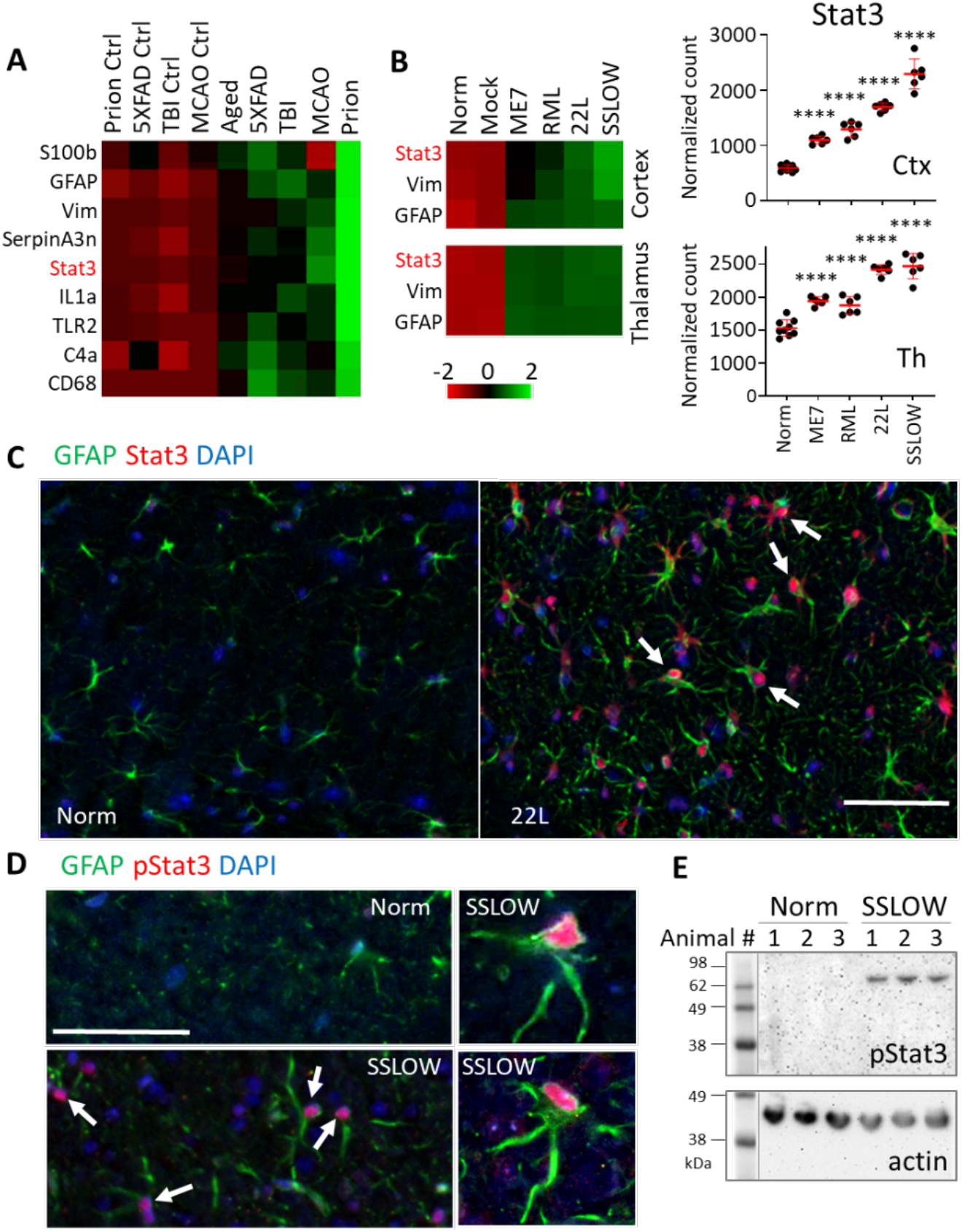
Stat3 and pStat3 are upregulated in prion-infected mice. A. Heat map showing expression of neuroinflammation-associated genes, including Stat3, in bulk brain tissue from 24-month-old C57BL/6J mice (aging), 10-month-old 5xFAD mice (Alzheimer’s disease model), C57BL/6J mice subjected to traumatic brain injury (TBI; 7 days post-injury), C57BL/6J mice subjected to middle cerebral artery occlusion (MCAO; analyzed 24 h after ischemia), SSLOW-infected C57BL/6J mice, and their corresponding control groups (n=3-9). B. Heat maps and corresponding quantification of Stat3, Vim, and GFAP expression in the cortex and thalamus of C57BL/6J mice infected IC with ME7, RML, 22L, or SSLOW prions and analyzed at the terminal stage of disease. Heat maps in **A** and **B** show log10- transformed normalized NanoString counts. Comparisons of different strains with non-infected control was performed by Brown-Forsythe and Welch’s ANOVA with Dunnett’s multiple comparisons test (Ctx) and ordinary one-way ANOVA with Dunnett’s T3 multiple comparisons test (Th), **** P<0.0001, n=6-9. C. Representative immunofluorescence images of brains from 22L-infected mice at the terminal stage of disease stained for GFAP and Stat3. D. Representative immunofluorescence images of brains from SSLOW-infected and age-matched non-infected C57BL/6J mice stained for GFAP and pStat3. Arrows indicate nuclear localization of Stat3 and pStat3. Scale bars 50 μm. E. Representative Western blot showing pStat3 expression in whole-brain homogenates from SSLOW-infected C57BL/6J mice collected at the terminal stage of disease.

All four mouse-adapted prion strains examined in this study (ME7, RML, 22L, and SSLOW) showed increased Stat3 expression (Fig. 1B). In both brain regions analyzed (cortex and thalamus), the magnitude of Stat3 upregulation followed the same pattern, with the strongest induction observed in SSLOW-infected mice (Fig. 1B). This ranking closely paralleled the degree of neuroinflammation previously reported for these prion strains [16].

Immunofluorescence staining of prion-infected brains demonstrated that Stat3, including phosphorylated Stat3 (pStat3), colocalized with GFAP-positive reactive astrocytes (Fig. 1C,D). Upon phosphorylation, Stat3 translocates to the nucleus, where it functions as a transcriptional regulator. Consistent with this mechanism, immunofluorescence revealed nuclear localization of pStat3 (Fig. 1D). Western blot analysis further confirmed a marked increase in pStat3 levels in prion-infected brains (Fig. 1E).

### Astrocyte-specific Stat3 deletion does not alter disease progression in SSLOW-infected mice

Among the four prion strains, SSLOW was selected for subsequent studies because it induces the strongest neuroinflammatory response and the greatest increase in Stat3 expression [16, 25, 61]. To determine whether astrocytic Stat3 signaling contributes to disease progression, we employed an astrocyte-specific conditional Stat3 knockout mouse model (Aldh1l1-CreERT-Stat3-floxP) [59]. In this model, Cre recombinase fused to a tamoxifen-inducible estrogen receptor (CreERT) is expressed under the control of the astrocyte-specific Aldh1l1 promoter. Administration of tamoxifen induces astrocyte-specific deletion of Stat3 in (Aldh1l1-CreERT)^+/-^Stat3-floxP)^+/+^ mice (Cre^+/-^). Cre-negative littermates (Cre^-/-^), which do not undergo recombination following tamoxifen treatment, served as controls. Wild-type C57BL/6J mice that did not receive tamoxifen were inoculated with SSLOW and included as a reference group.

The effect of astrocyte-specific Stat3 deletion was evaluated using two experimental formats: (i) mice were inoculated intracerebrally (IC) with SSLOW and treated with tamoxifen at the onset of clinical disease (14 weeks post-inoculation; SSLOW+Tam), or (ii) mice were treated with tamoxifen before IC inoculation with SSLOW (Tam+SSLOW) (Fig. 2A).

**Figure 2.**
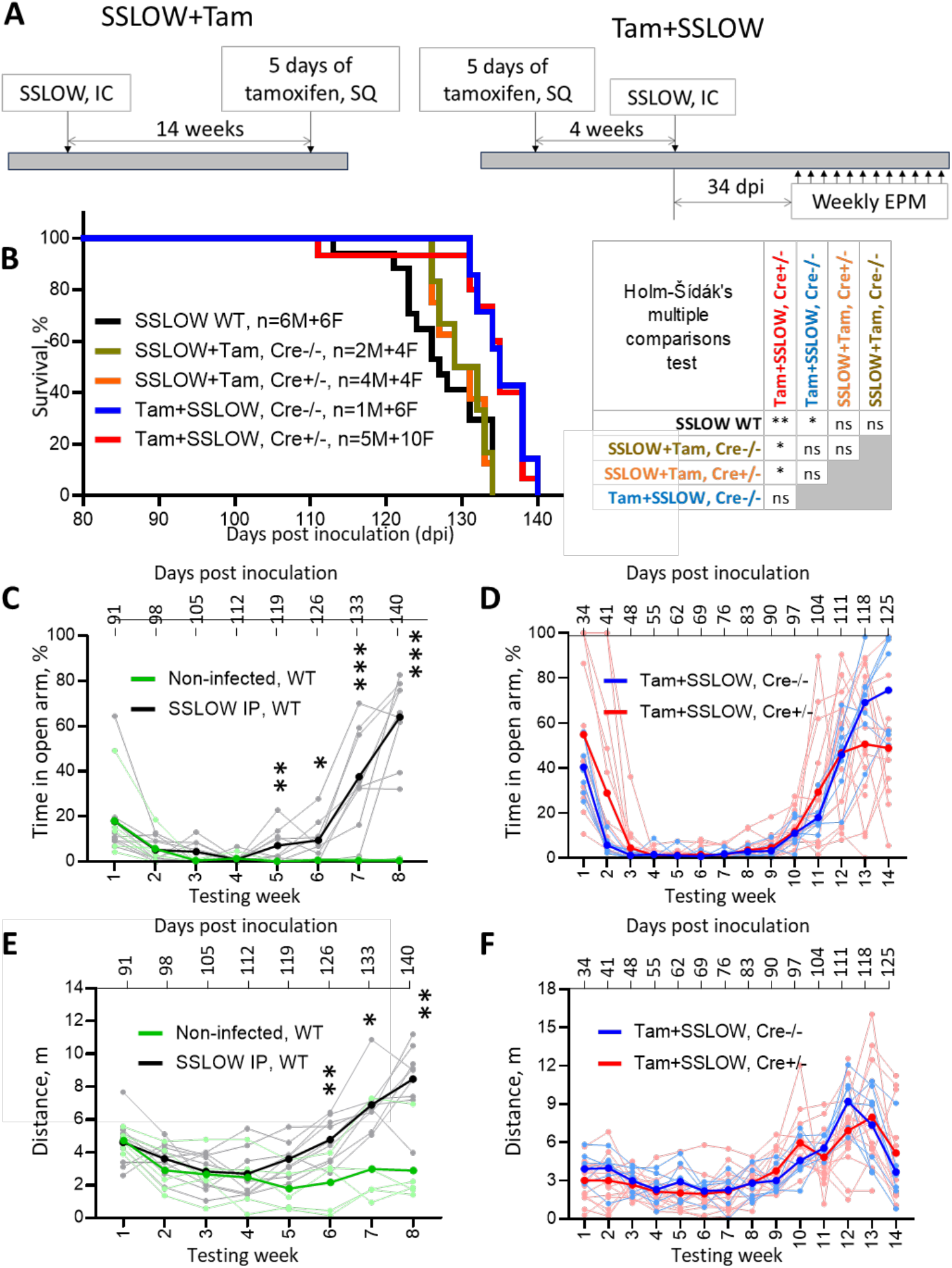
Astrocyte-specific Stat3 deletion does not alter disease progression in SSLOW-infected mice. A. Experimental design of the SSLOW+Tam and Tam+SSLOW formats. B. Kaplan-Meier survival curves for Cre^+/-^ and Cre^-/-^ mice in the SSLOW+Tam and Tam+SSLOW experiments and wild-type C57BL/6J mice (WT) infected via IC route. WT mice did not receive tamoxifen. Statistical significance was determined using the Mantel-Cox log-rank test with Holm-Šídák’s multiple comparisons test. C,E. An example of elevated plus maze (EPM) analysis of wild-type C57BL/6J mice infected with SSLOW via IP route and age-matched non-infected controls. D,F. EPM analysis of Cre^+/-^ and Cre^-/-^ mice in the Tam+SSLOW experiment. Shown are the percentage of time spent in the open arms (**C, D**) and total distance traveled during each session (**E, F**). Mice were tested weekly beginning at the preclinical stage. Thick lines indicate group means and thin lines represent individual animals. *P*<0.05, \**P*<0.01, \*\**P*<0.001; statistical analysis of Cre^-/-^ and Cre^+/-^ groups for each time point were performed as described in Methods.

In both experimental formats, Cre^+/-^ and Cre^-/-^ mice reached the terminal stage of disease at comparable times, indicating that astrocyte-specific Stat3 deletion did not significantly alter disease progression or survival following SSLOW infection (Fig. 2B). However, both tamoxifen-treated groups survived longer than SSLOW-infected wild-type mice that did not receive tamoxifen (Fig. 2B). This effect was observed in both experimental paradigms and reached statistical significance when tamoxifen was administered before prion inoculation (Tam+SSLOW). These findings suggest that tamoxifen treatment itself contributed to prolonged survival, with a greater protective effect when administered prior to infection.

To complement conventional clinical scoring, behavioral changes were monitored in the Tam+SSLOW cohort using the elevated plus maze (EPM). Beginning at the preclinical stage, mice were tested weekly. Following one or two habituation sessions, healthy mice developed a strong preference for the closed arms of the maze, which persisted throughout the observation period (Fig. 2C). As prion disease progressed, infected mice gradually lost this preference and spent increasing amounts of time in the open arms (Fig. 2C,D). Disease progression was also accompanied by increased locomotor activity, reflected by longer distances traveled during the test, consistent with hyperactivity. This increase persisted until the terminal stage, when locomotor activity declined because of progressive motor impairment (Fig. 2E,F). In our experience, the EPM provides a sensitive and objective measure of behavioral changes during prion disease that complements conventional clinical scoring. Nevertheless, EPM analysis detected no differences in the temporal progression of behavioral abnormalities between Cre^+/-^ and Cre^-/-^ mice (Fig. 2D,F).

### Astrocyte-specific Stat3 deletion delays disease progression in 22L-infected male mice

We next performed analogous experiments using the 22L prion strain, which is characterized by a longer incubation period and a milder neuroinflammatory response than SSLOW. Similarly to the SSLOW experiments, mice were treated with tamoxifen either following inoculation with prions (10 weeks post-inoculation; 22L+Tam) or before prion inoculation (Tam+22L) (Fig. 3A). As with the SSLOW model, both tamoxifen-treated Cre^+/-^ and Cre^-/-^ mice reached the terminal stage of disease later than wild-type mice that did not receive tamoxifen (Fig. 3B). This effect was more pronounced in the Tam+22L format than in the 22L+Tam format, consistent with the trend observed following SSLOW infection.

**Figure 3.**
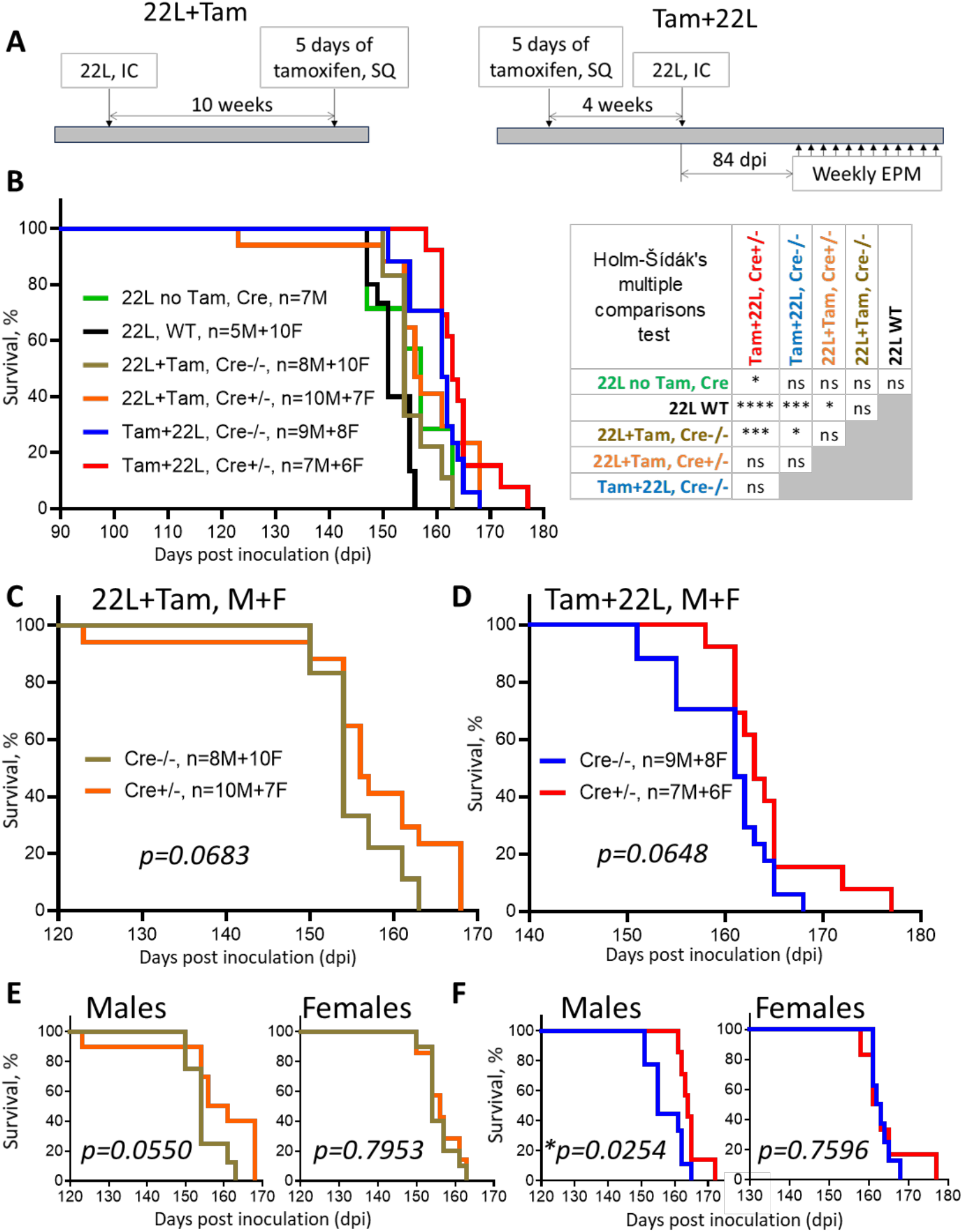
Astrocyte-specific Stat3 deletion delays disease progression in 22L-infected male mice. A. Experimental design of the 22L+Tam and Tam+22L formats. B. Kaplan-Meier survival curves for Cre^+/-^ and Cre^-/-^ mice in the 22L+Tam and Tam+22L experiments, wild-type 22L-infected C57BL/6J mice, and untreated 22L-infected Cre controls (combined Cre^+/-^ and Cre^-/-^ mice). All groups were inoculated via IC route. C,D. Kaplan-Meier survival curves comparing Cre^+/-^ and Cre^-/-^ mice in the 22L+Tam (**C**) and Tam+22L (**D**) formats. E,F. Sex-stratified Kaplan-Meier survival curves comparing Cre^+/-^ and Cre^-/-^ mice in the 22L+Tam (**E**) and Tam+22L (**F**) formats. Statistical significance was determined using the Mantel- Cox log-rank test (B-F), followed by Holm-Šídák’s multiple comparisons test when needed (B).

To determine whether this delay truly reflected the effect of tamoxifen, we included an additional control group consisting of Cre^+/-^ and Cre^-/-^ mice inoculated with 22L but not treated with tamoxifen (designated Cre) (Fig. 3B). Disease progression in the untreated Cre group was closer to the WT. Also, mice treated with tamoxifen before prion inoculation (Tam+22L) showed longer incubation time than either the untreated Cre group or the 22L+Tam Cre^-/-^ group, suggesting that early tamoxifen administration confers a protective effect.

Within both experimental formats, Cre^+/-^ mice survived slightly longer than Cre^-/-^ littermates, although the differences reached only borderline statistical significance (Fig. 3C,D). Sex-stratified analysis revealed that this effect was largely restricted to males. In both experimental formats, male Cre^+/-^ mice exhibited a more pronounced extension of survival than male Cre^-/-^ mice, with the greatest difference observed in the Tam+22L format (Fig. 3E,F). In contrast, female Cre^+/-^ and Cre^-/-^ mice showed virtually identical survival curves in both treatment formats.

Behavioral analysis using EPM was consistent with the survival data. Male Cre^+/-^ mice in the Tam+22L cohort displayed a delayed onset of behavioral abnormalities compared with Cre^-/-^ littermates (Fig. 4A,B), whereas female mice exhibited indistinguishable disease trajectories regardless of genotype (Fig. 4C,D). Together, these findings demonstrate that astrocyte-specific Stat3 deletion produces a modest delay in disease progression in the 22L model, an effect that is most evident when recombination is induced before prion infection and is largely confined to male mice.

**Figure 4.**
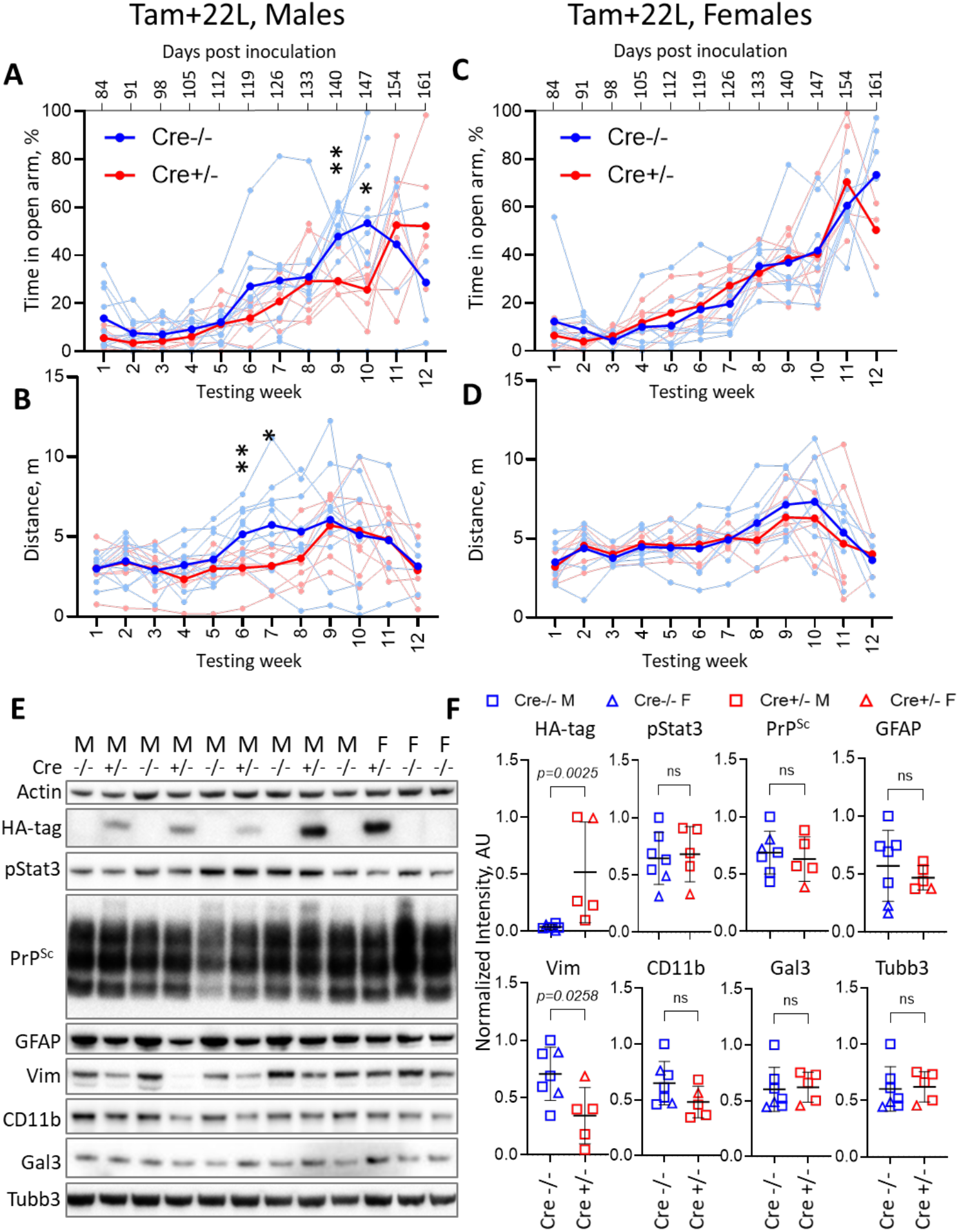
Behavioral analysis and neuropathological characterization of the Tam+22L groups. A-D. EPM analysis of Cre^+/^- and Cre^-/-^ mice in the Tam+22L experiment stratified by sex. Male mice are shown in **A** and **B**, and female mice in **C** and **D**. Percentage of time spent in the open arms (**A, C**) and total distance traveled during each session (**B, D**) are shown. Mice were tested weekly beginning at the preclinical stage. Thick lines indicate group means and thin lines represent individual animals. \**P*<0.05, \*\**P*<0.01; statistical analysis of Cre^-/-^ and Cre^+/-^ groups for each time point were performed as described in Methods. E. Representative Western blots of whole-brain homogenates collected at the terminal stage of disease from Cre^+/-^ (n=5) and Cre^-/-^ (n=7) mice in the Tam+22L experiment. F. Quantification of HA-tag, pStat3, PrP^Sc^, GFAP, vimentin (Vim), CD11b, Gal3, and βIII-tubulin (Tubb3) levels shown in **E**. Protein levels were normalized to β-actin. Squares and triangles represent male and female mice, respectively. Statistical analysis of Cre^-/-^ (n=7) and Cre^+/-^ (n=5) groups were performed as described in Methods. ns, not significant.

The 22L experiments therefore revealed a mild but statistically significant protective effect of astrocyte-specific Stat3 deletion that was not observed in the SSLOW model. The greater effect in the Tam+22L format suggests that Stat3 contributes primarily during the early establishment of astrocyte reactivity rather than after reactive astrocytes have already developed. The apparent sex specificity further indicates that the impact of astrocytic Stat3 signaling on prion disease progression may be influenced by biological sex. Notably, male mice were underrepresented in the SSLOW cohorts, which consisted predominantly of females. Consequently, the absence of a detectable phenotype in the SSLOW experiments may reflect both the more severe inflammatory environment elicited by this strain and differences in the sex composition of the experimental groups.

### Astrocyte-specific Stat3 deletion attenuates astrocyte reactivity

To determine whether the modest delay in disease progression observed in 22L-infected Cre^+/-^ mice was accompanied by changes in neuropathology, brains collected at the terminal stage of the Tam+22L experiment were analyzed by Western blot. PrP^Sc^ accumulation was comparable between Cre^+/-^ and Cre^-/-^ mice, indicating that astrocyte-specific Stat3 deletion did not alter prion accumulation (Fig. 4E,F). Likewise, no differences in total pStat3 levels were detected between the two genotypes (Fig. 4E,F). Although unexpected, this finding can be explained by the cell type specificity of the knockout, which was restricted to Aldh1l1-positive astrocytes. Other brain cell types, including microglia, neurons, oligodendrocytes, and endothelial cells, express comparable levels of Stat3 [62], likely masking changes in astrocytic pStat3 in whole-brain lysates.

Both Cre^+/-^ and Cre^-/-^ mice also carried the tamoxifen-inducible HA-RiboTag allele, enabling verification of Cre-mediated recombination by HA-tag expression. As expected, robust HA expression was detected only in tamoxifen-treated Cre^+/-^ mice (Fig. 4E,F). Notably, HA expression varied among individual Cre^+/-^ mice, suggesting variability in the efficiency of Cre-mediated recombination and, consequently, Stat3 deletion.

At the terminal stage of disease, astrocyte-specific Stat3 deletion did not measurably alter GFAP expression. In contrast, the astrocyte reactivity marker vimentin (Vim) was significantly reduced in Cre^+/-^ mice relative to Cre^-/-^ controls (Fig. 4E,F). Astrocyte-specific Stat3 deletion also did not affect bulk neuronal count or microglial reactivity, as assessed by Western blot analysis of the neuronal marker βIII- tubulin (Tubb3) and the microglial markers CD11b and galectin-3 (Gal3).

### Astrocyte-specific Stat3 deletion delays the development of astrocyte reactivity

To determine whether the effects of astrocyte-specific Stat3 deletion were more pronounced during the early stages of disease, the Tam+22L experiment was repeated, and brains were collected at the late preclinical stage (91 dpi) and at the onset of clinical disease (111 dpi). As in the terminal-stage analysis, successful Cre-mediated recombination in Cre^+/-^ mice was confirmed by HA-tag expression on Western blots (Fig. 5A-D). Consistent with the terminal-stage findings, total pStat3 levels and PrP^Sc^ accumulation in whole-brain lysates were comparable between Cre^+/-^ and Cre^-/-^ mice at both time points (Fig. 5A-D). Similar to the terminal-stage analysis, the lack of change in bulk pStat3 levels likely reflects the astrocyte-specific nature of the knockout, as Stat3 is also abundantly expressed by other brain cell types [62]. Interestingly, βIII-tubulin (Tubb3) levels were modestly higher in Cre^+/-^ mice at both stages, suggesting a potential neuroprotective effect of astrocyte-specific Stat3 deletion during the late preclinical and early clinical phases of prion disease (Fig. 5A-D).

**Figure 5.**
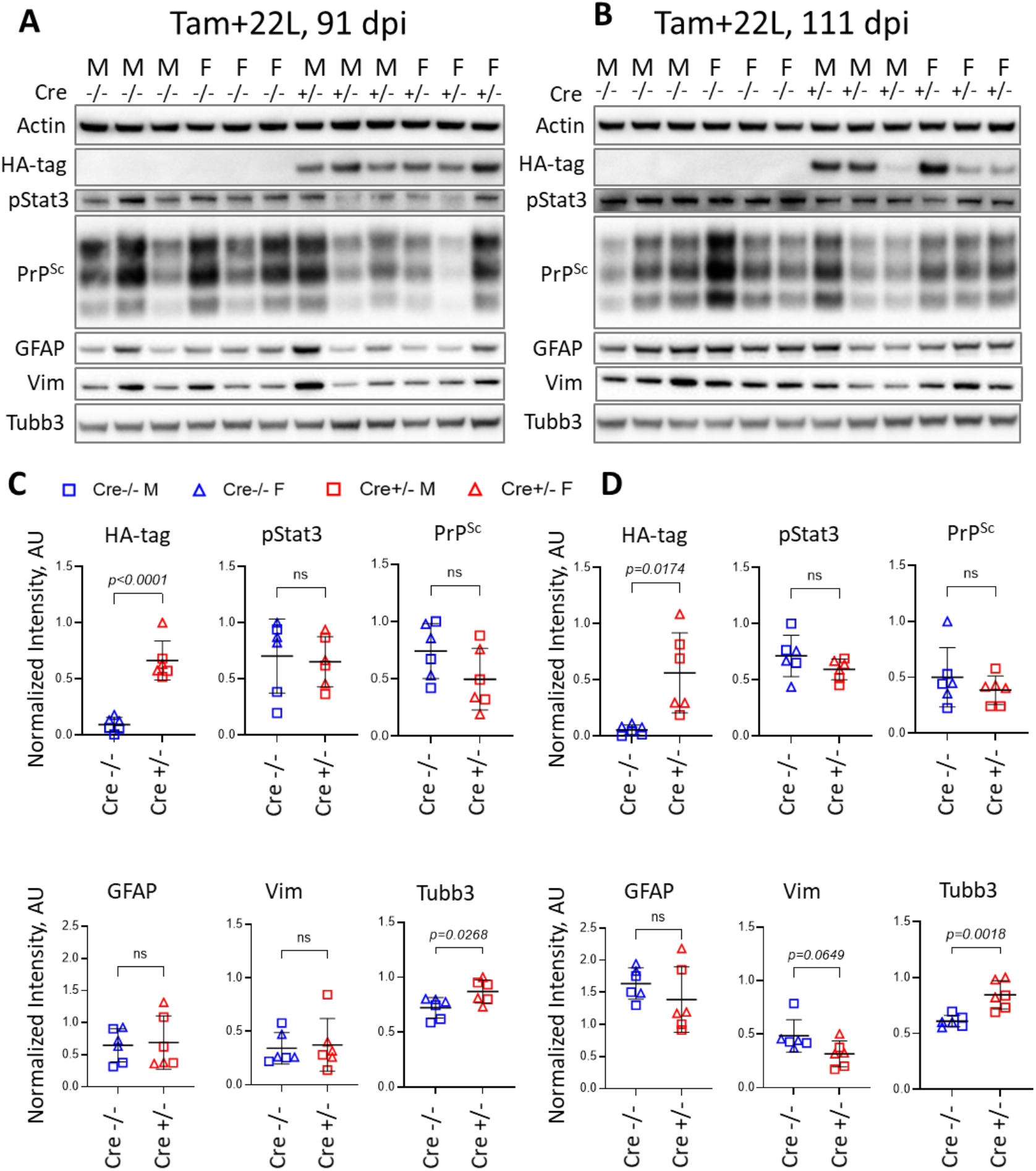
Analysis of brain pathology during the early stages of disease in the Tam+22L experiment. A,B. Representative Western blots of whole-brain homogenates from Cre^+/−^ and Cre^−/−^ mice in the Tam+22L experiment collected at 91 dpi (**A**) and 111 dpi (**B**). C,D. Quantification of HA-tag, pStat3, PrP^Sc^, GFAP, Vim, and βIII-tubulin (Tubb3) levels in whole-brain homogenates from Cre^+/−^ and Cre^−/−^ mice analyzed at 91 dpi (**C**) and 111 dpi (**D**). Protein levels were normalized to actin. Squares and triangles represent male and female mice, respectively. Statistical analysis of Cre^-/-^ (n=6) and Cre^+/-^ (n=6) groups were performed as described in Methods. ns, not significant.

Western blot analysis revealed comparable GFAP levels in whole-brain lysates from Cre^+/-^ and Cre^-/-^ mice at both time points (Fig. 5A-D). In contrast, vimentin (Vim) levels showed a downward trend in Cre^+/-^ mice at 111 dpi, consistent with the significant reduction observed at the terminal stage (Fig. 5A-D).

Because Western blot analysis of whole-brain homogenates may obscure region-specific changes in astrocyte reactivity, GFAP expression was next examined by immunofluorescence (Fig. 6A). Quantification of cortical GFAP immunoreactivity revealed significantly lower GFAP levels in Cre^+/-^ mice than in Cre^-/-^ controls at 91 dpi, whereas no differences were detected at 111 dpi (Fig. 6B). In contrast, cortical IBA1 immunoreactivity was comparable between the two genotypes at both time points (Fig. 6B), indicating that astrocyte-specific Stat3 deletion did not measurably alter microglial activation. Together, these findings suggest that astrocyte-specific Stat3 deletion delays the development of astrocyte reactivity but does not prevent its progression at later stages of prion disease.

**Figure 6.**
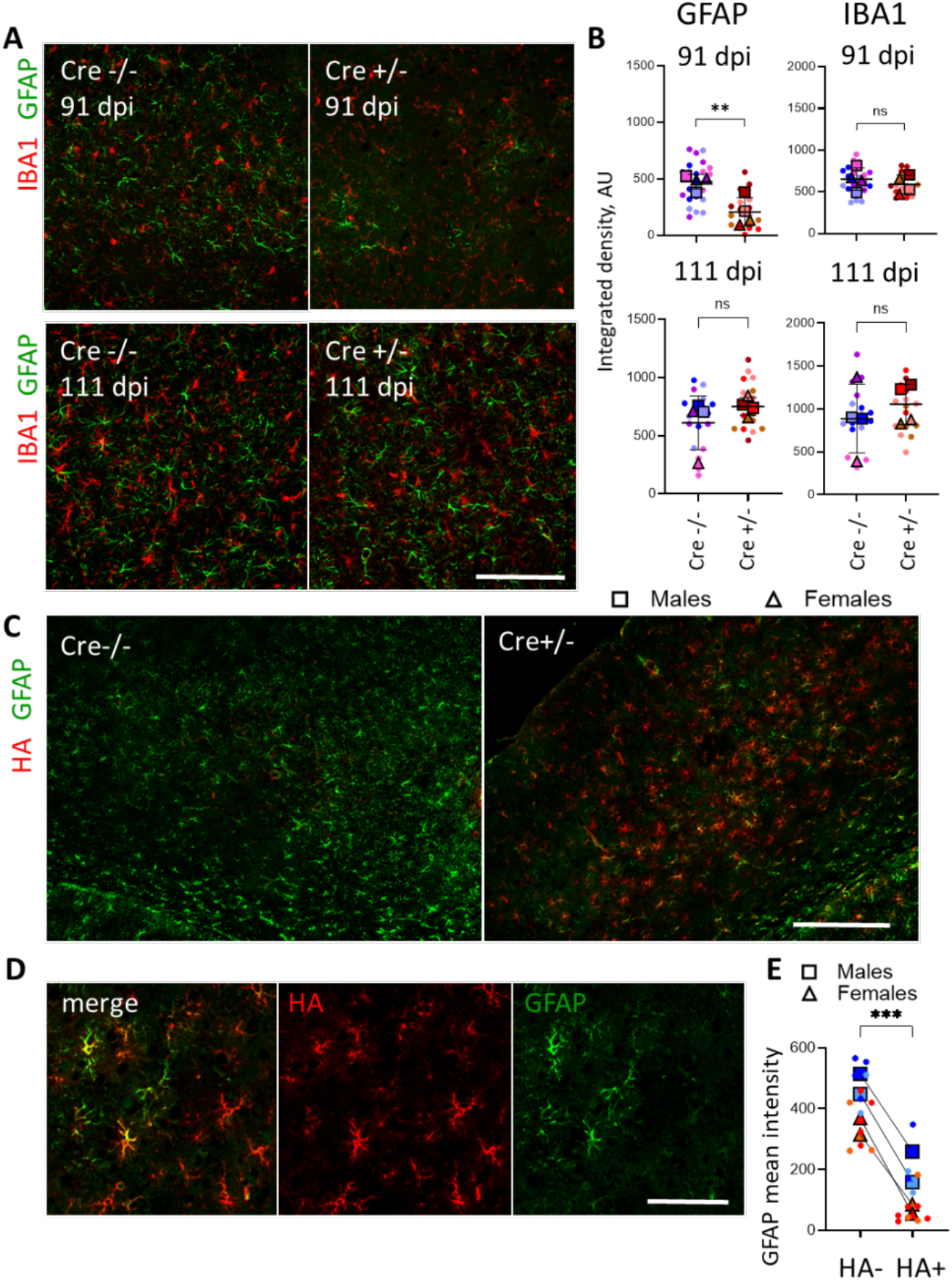
Astrocyte-specific Stat3 deletion delays the development of astrocyte reactivity. A. Representative immunofluorescence images of the cortex from Cre^−/−^ and Cre^+/−^ mice in the Tam+22L experiment co-immunostained for GFAP and IBA1 and analyzed at 91 and 111 dpi. B. Quantification of cortical GFAP and IBA1 immunoreactivity shown in (**A**). Integrated fluorescence intensity was measured for each marker. Superplot colors represent individual brains; dots represent individual fields of view; average values for each brain are shown as squares (males) and triangles (females); means ± SD are marked by black lines. Comparisons of means for Cre^-/-^ (n=4) and Cre^+/-^ (n=4) groups were performed as described in Methods. \*\**P*<0.01, ns, not significant. C. Representative cortical sections from Cre^−/−^ and Cre^+/−^ mice from the Tam+22L experiment collected at 91 dpi and co-immunostained for GFAP and the HA epitope. HA-positive astrocytes were detected only in Cre^+/−^ mice, confirming successful Cre-mediated recombination. D. Representative fluorescent images illustrating HA-positive and HA-negative astrocytes in the cortex of Cre^+/−^ mice. D. Quantification of GFAP mean fluorescence intensity in HA-positive and HA-negative astrocytes from the cortex of Cre^+/−^ mice at 91 dpi. Superplot colors represent individual brains; dots represent individual fields of view; average values for each brain are shown as squares (males) and triangles (females). N = 4 mice. ***P < 0.001, by paired *t*-test. Scale bars 100 μm in A and D, and 200 μm in C.

To determine whether Cre-mediated recombination reduced astrocyte reactivity at the single-cell level, brain sections collected at 91 dpi were co-immunostained for HA and GFAP. Numerous HA-positive cells were detected in Cre^+/-^ mice but were absent from Cre^-/-^ controls, confirming selective recombination in Cre^+/-^ animals (Fig. 6C,D). Quantitative analysis further demonstrated that HA-positive astrocytes expressed significantly lower levels of GFAP than neighboring HA-negative astrocytes within the same brains (Fig. 6D,E). Reduced GFAP expression in HA-positive cells was observed in both male and female mice (Fig. 6E). These findings demonstrate that Cre-mediated recombination occurred in only a subset of astrocytes and that recombined astrocytes exhibited a lower degree of reactive transformation than unrecombined neighboring cells.

### Tamoxifen-induced recombination results in partial astrocyte-specific Stat3 deletion

Because astrocyte-specific Stat3 deletion produced only a modest effect on prion disease progression, we next assessed the efficiency of Cre-mediated recombination using HA expression as a surrogate marker. In brains from the Tam+22L cohort, tamoxifen-treated Cre^+/-^ mice exhibited robust HA immunoreactivity throughout all brain regions examined (Fig. 7A). However, quantification of brain sections coimmunostained for GFAP and HA revealed that the proportion of HA-positive astrocytes varied among brain regions, ranging from approximately 40% to 70% (Fig. 7B-D). These findings indicate that the tamoxifen-inducible CreERT system achieved only partial astrocyte-specific Stat3 deletion and that the efficiency of recombination was region dependent.

**Figure 7.**
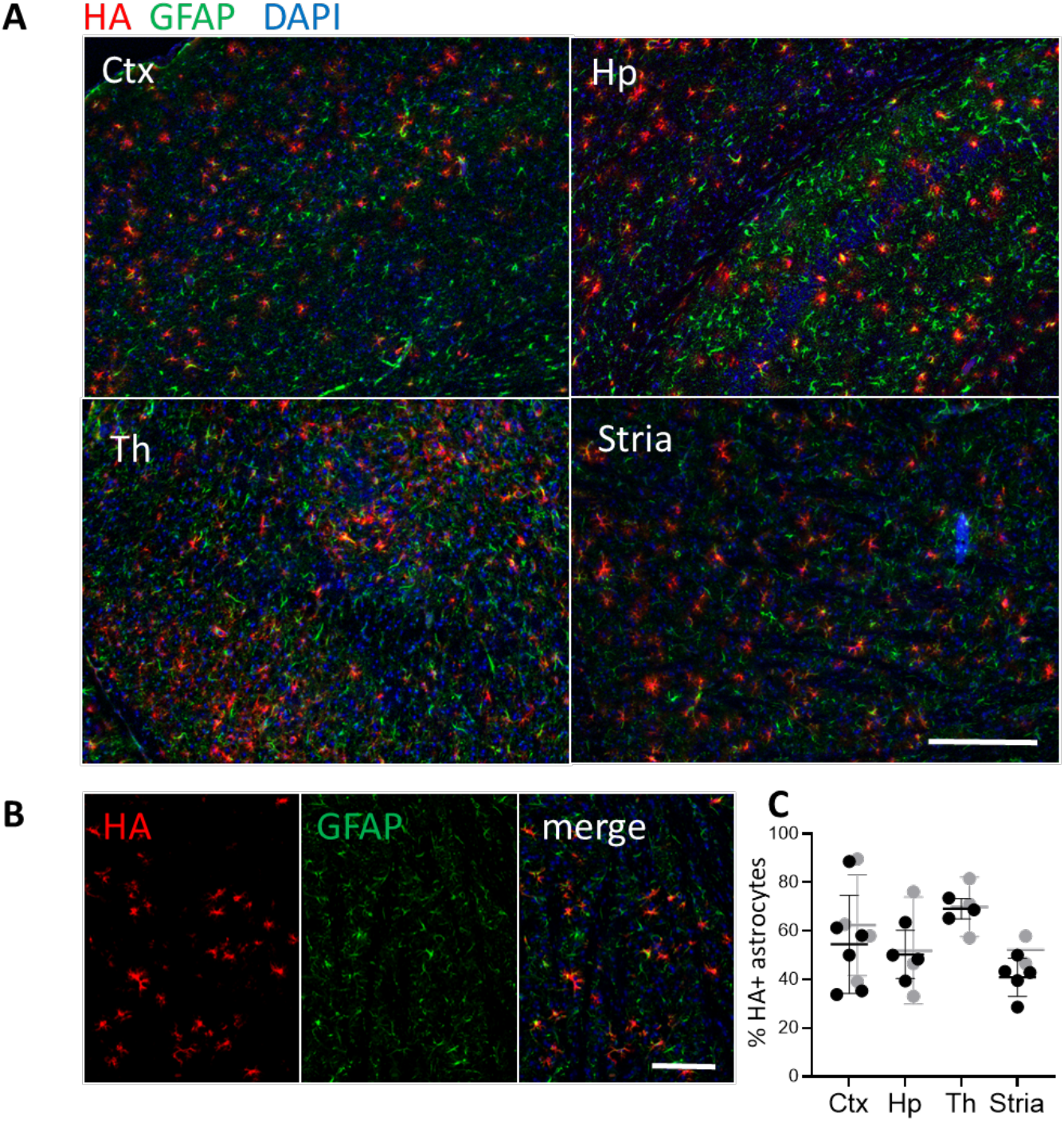
Tamoxifen-induced recombination results in partial astrocyte-specific Stat3 deletion. A. Representative immunofluorescence images of the cortex (Ctx), hippocampus (Hp), thalamus (Th), and striatum (Stria) from Cre^+/−^ mice in the Tam+22L experiment, collected at the terminal stage and co- immunostained for GFAP and the HA epitope. B. Representative images of the striatum showing the GFAP and HA channels separately and as a merged image, illustrating astrocytes that underwent Cre-mediated recombination. Scale bars 200 μm. C. Quantification of the percentage of HA-positive astrocytes in the cortex, hippocampus, thalamus, and striatum of Cre^+/−^ mice from the Tam+22L experiment at the terminal stage. Black and gray symbols represent individual mice, dots indicate individual fields of view, and lines represent the mean value for each mouse ±SD.

## Discussion

Stat3 signaling is considered as the core master regulator of astrocyte reactivity in acute and chronic conditions [40–42]. By selectively depleting Stat3 in astrocytes, our study sought to determine whether reactive astrogliosis serves a protective function, contributes to disease pathogenesis, or represents an epiphenomenon of chronic neurodegeneration.

A central message emerging from this study is that although Stat3 signaling is one of the most robust molecular signatures of reactive astrocytes in prion disease, it is not a major determinant of disease progression. Instead, our results suggest that astrocytic Stat3 primarily regulates the magnitude and timing of astrocyte reactivity, while exerting only a modest influence on neurodegeneration. This conclusion is supported by several key findings: (i) Stat3 was strongly induced in reactive astrocytes across multiple prion strains; (ii) astrocyte-specific Stat3 deletion consistently reduced astrocyte activation, particularly during early disease stages; yet (iii) the overall impact on incubation time was limited, becoming detectable only in male mice infected with the less inflammatory 22L strain and primarily when Stat3 deletion preceded prion infection.

Among several neurological conditions associated with neuroinflammation examined in the current work, prion disease displayed the strongest Stat3 induction, exceeding that observed during aging, Alzheimer’s disease, traumatic brain injury or ischemia (Fig. 1A). Moreover, all four mouse-adapted prion strains tested exhibited increased Stat3 expression (Fig. 1B), with the magnitude of induction closely paralleling the severity of neuroinflammation previously reported for these strains [16]. Despite this prominent activation, astrocyte-specific Stat3 deletion had little effect on disease progression. SSLOW- infected mice, which develop the strongest neuroinflammatory response and exhibit the highest Stat3 induction [16, 25, 61], showed no detectable benefit from Stat3 deletion. In contrast, only a mild extension of survival was observed in 22L-infected animals, although only in males, with the greatest benefit occurring when Stat3 deletion was induced before prion inoculation. These findings suggest that astrocytic Stat3 contributes primarily during the establishment of reactive astrocyte programs rather than during advanced disease. Once extensive gliosis and neurodegeneration have developed, multiple parallel inflammatory pathways may compensate for loss of Stat3 signaling [63], thereby limiting its overall contribution to disease progression.

Increasing evidence indicates that microglia are major upstream regulators of astrocyte reactivity during neurodegeneration [21, 35, 54]. Upon activation, reactive microglia establish and maintain a pro-inflammatory environment through the secretion of multiple cytokines, including IL-1α, TNF-α, and the complement component C1q. These factors act synergistically to induce the transition of homeostatic astrocytes into a reactive phenotype, including the neurotoxic astrocyte [35]. In prion disease, reactive microglia have likewise been shown to drive the acquisition of neurotoxic astrocyte phenotypes, highlighting extensive crosstalk between these two glial populations [54, 55]. Within this framework, the robust activation of Stat3 observed in astrocytes during prion disease may represent a downstream consequence of persistent microglia-derived inflammatory signaling rather than the initiating event in astrocyte activation. This model is consistent with our finding that astrocyte-specific Stat3 deletion attenuated astrocyte reactivity but had little effect on overall microglial activation or disease progression, suggesting that Stat3 functions downstream of inflammatory cues generated by reactive microglia. The persistence of microglial activation in Stat3-deficient mice may therefore provide sufficient pro-inflammatory signaling to sustain astrocyte activation through compensatory pathways [63, 64], thereby limiting the impact of astrocytic Stat3 depletion on disease outcome.

Our histological analyses demonstrate that Stat3 is an important regulator of astrocyte activation. Whole-brain Western blot analysis revealed a downward trend in the astrocyte reactivity marker Vim beginning at clinical onset, with a significant reduction at the terminal stage. In contrast, regional immunofluorescence detected significantly lower cortical GFAP expression in Stat3-deficient mice during the late preclinical stage, whereas this difference was no longer evident at disease onset. Importantly, single-cell analysis demonstrated that recombined HA-positive astrocytes consistently exhibited lower GFAP expression than neighboring unrecombined astrocytes within the same brain, providing direct evidence for a cell-autonomous effect of Stat3 depletion. The distinct temporal patterns observed for GFAP and Vim suggest that Stat3 differentially regulates astrocyte subpopulations expressing these markers. This interpretation is consistent with previous studies demonstrating considerable heterogeneity among reactive astrocytes in chronic neurodegenerative diseases and showing that, although GFAP and Vim are frequently co-expressed, they do not define identical astrocyte populations [16, 23, 34, 65]. Collectively, these findings indicate that Stat3 directly promotes astrocyte reactivity in a cell-autonomous manner. However, astrocyte activation ultimately progressed despite Stat3 deletion, suggesting that Stat3 facilitates the initiation and amplification of reactive astrocyte programs but is not absolutely required for their eventual development. Importantly, attenuation of astrocyte reactivity occurred without detectable changes in PrP^Sc^ accumulation or microglial activation (Fig. 4). Neither biochemical analysis of PrP^Sc^ nor Western blotting for markers of reactive microglia CD11b and Gal3 revealed significant differences between control and Stat3-deficient animals. Notably, Gal3 has emerged as a particularly sensitive marker of perturbations in microglial reactivity [17, 25, 26, 66]. Its robust induction during prion disease makes the absence of changes in Gal3 expression between Cre^+/-^ and Cre^-/-^ animals especially informative. Similarly, cortical IBA1 immunoreactivity remained unchanged in Cre^+/-^ animals. These findings indicate that astrocytic Stat3 signaling is dispensable for prion replication and does not substantially alter the overall microglial response. Instead, Stat3 appears to regulate intrinsic astrocyte activation independently of the core mechanisms governing prion propagation or microglial activation.

The modest increase in neuronal βIII-tubulin observed during preclinical and early clinical stages raises the intriguing possibility that reduced astrocyte reactivity provides transient neuroprotection. Although this effect was insufficient to substantially delay disease progression, it suggests that reactive astrocytes regulated by Stat3 may contribute to neuronal dysfunction. Similar neuroprotective effects following astrocytic Stat3 deletion have been reported in Alzheimer’s disease mouse models [46–48], where excessive astrocyte activation can impair neuronal homeostasis through altered metabolic support, inflammatory mediator release, or disruption of synaptic function.

An unexpected finding was the influence of tamoxifen treatment itself on disease progression. Both SSLOW and 22L experiments demonstrated prolonged survival in tamoxifen-treated mice relative to untreated wild-type controls, particularly when tamoxifen administration preceded prion inoculation (Fig. 2,3). Tamoxifen possesses well-documented anti-inflammatory and neuroprotective properties independent of Cre-mediated recombination [67–73], and these effects likely contributed to delayed disease onset in our experiments. This observation emphasizes the importance of including tamoxifen-treated Cre-negative controls when using inducible Cre systems in neurodegenerative disease models and suggests that tamoxifen may influence prion pathogenesis through mechanisms unrelated to Stat3 deletion.

Our study also revealed an unexpected sex-specific effect, with protection observed only in male mice. Sex-dependent differences in astrocyte biology and neuroimmune responses have become increasingly recognized across multiple neurological disorders and aging [74–76]. Astrocytes exhibit sexually dimorphic transcriptional profiles, respond differently to inflammatory stimuli, and are influenced by sex hormones that can directly modulate Stat3 signaling pathways [77–80]. Although our study was not designed to dissect these mechanisms, the preferential benefit observed in males suggests that astrocytic Stat3 signaling may interact with sex-dependent regulatory pathways during prion disease. Future studies using balanced cohorts and mechanistic analyses will be necessary to determine the basis of this sexual dimorphism.

A major limitation of the present study is the incomplete efficiency of tamoxifen-induced recombination. Depending on brain region, approximately 40-70% of astrocytes underwent recombination, indicating that a substantial proportion of astrocytes retained intact Stat3 expression (Fig. 7). This mosaic deletion likely reduced the overall impact of the knockout and may explain why total brain pStat3 levels remained unchanged despite successful recombination in individual astrocytes. Moreover, it is possible that neighboring Stat3-intact astrocytes could partially compensate for Stat3-deficient cells through intercellular signaling, thereby preserving the overall inflammatory environment. Consequently, the relatively modest phenotypic effects observed here should probably be considered a conservative estimate of the contribution of astrocytic Stat3 signaling. Future studies employing constitutive astrocyte-specific deletion, viral gene delivery, or more efficient inducible recombination strategies will be required to fully define the role of astrocytic Stat3 during prion disease.

Finally, our findings contribute to the broader discussion regarding the functional significance of reactive astrocytes in neurodegeneration. Reactive astrocytes are often viewed as major drivers of neuronal dysfunction [35, 63, 81], and Stat3 has emerged as one of the principal regulators of astrocyte reactivity [40–42]. However, our data suggest that reducing astrocyte reactivity alone produces only limited therapeutic benefit in prion disease. Rather than serving as a primary pathogenic driver, Stat3-dependent astrocyte activation may represent one component of a highly interconnected multicellular response involving microglia, neurons, endothelial cells and infiltrating immune signals [82–84]. Consequently, therapeutic strategies directed solely at astrocytic Stat3 are unlikely to substantially modify disease progression. More effective approaches may require simultaneous targeting of multiple inflammatory pathways or earlier intervention before the establishment of irreversible neurodegenerative cascades.

In conclusion, we demonstrate that Stat3 is robustly activated in reactive astrocytes across multiple prion strains and serves as an important regulator of astrocyte activation. Nevertheless, astrocyte-specific Stat3 deletion only modestly delays prion disease progression in 22L-infected males, while reducing astrocyte reactivity without affecting prion accumulation or global microglial activation. These findings indicate that astrocytic Stat3 is required for full development of reactive astrocytes but is not a dominant driver of prion pathogenesis, highlighting the remarkable redundancy of inflammatory signaling networks operating during chronic prion-induced neurodegeneration.

## List of abbreviations

22L: mouse adapted prion strain
Aldh1l1: aldehyde dehydrogenase 1 family member L1
BBB: blood brain barrier
BH: brain homogenates
CD11b: Integrin subunit alpha M
CNS: Central Nervous System
Dlg4: Discs Large Homolog 4, also known as PSD95
Dpi: days post inoculation
EPM: elevated plus maze
Gal3: Galectin-3
GFAP: Glial fibrillary acidic protein
HA: Hemagglutinin
Jak2: Janus Kinase 2
IL-6: Interleukin-6
IC: intracranial route
ME7: mouse adapted prion strain
PrP^C^: normal, cellular isoform of the prion protein
PrP^Sc^: infectious, disease-associated, pathogenic form of the prion protein
pStat3: phosphorylated Signal Transducer and Activator of Transcription 3
RML: mouse adapted prion strain
SSLOW: mouse adapted prion strain
Stat3: Signal Transducer and Activator of Transcription 3
Tam: tamoxifen
Vim: vimentin

## Declarations

### Ethics approval and consent to participate

The study was carried out in strict accordance with the recommendations in the Guide for the Care and Use of Laboratory Animals of the National Institutes of Health. The animal protocol was approved by the Institutional Animal Care and Use Committee of the University of Maryland, Baltimore (Assurance Number: A32000-01; Protocol Numbers: 1120001 and 00000166-1).

### Consent for publication

Not applicable

### Availability of data and materials

All data generated or analyzed during this study are included in this published article and its supplementary information file.

### Authors’ contributions

NM and IB designed the study; NM, NP, OM, TS and OB performed experiments; KM performed animal procedures, monitored behavioral changes and scored the disease signs; NM, NP and TS analyzed the data; IB and NM wrote the manuscript. All authors read, edited and approved the final manuscript.

## Acknowledgments

We are grateful to Michael Sofroniew for his generous gift of Aldh1l1-CreERT-Stat3-floxP mouse model.

